# A hierarchical approach to model decision making: a study in chemotactic behavior of *Escherichia coli*

**DOI:** 10.1101/569277

**Authors:** Safar Vafadar, Kaveh Kavousi, Hadiseh Safdari, Ata Kalirad, Mehdi Sadeghi

**Affiliations:** Laboratory of Biological Complex Systems and Bioinformatics (CBB), Institute of Biochemistry and Biophysics, University of Tehran, Tehran, Iran; School of Biological Science, Institute for Research in Fundamental Sciences (IPM), Tehran, Iran; National Institute of Genetic Engineering and Biotechnology (NIGEB), Tehran, Iran

## Abstract

Reducing the complex behavior of living entities to its underlying physical and chemical processes is a formidable task in biology. Complex behaviors can be characterized as decision making: the ability to process the incoming information via an intracellular network and act upon this information to choose appropriate strategies. Motility is one such behavior that has been the focus many modeling efforts in the past. Our aim is to reduce the chemotactic behavior in E. coli to its molecular constituents in order to paint a comprehensive and end-to-end picture of this intricate behavior. We utilize a hierarchical approach, consisting of three layers, to achieve this goal: at the first level, chemical reactions involved in chemotaxis are simulated. In the second level, the chemical reactions give rise to the mechanical movement of six independent flagella. At the last layer, the two lower layers are combined to allow a digital bacterium to receive information from its environment and swim through it with verve. Our results are in concert with the experimental studies concerning the motility of E. coli cells. In addition, we show that our detailed model of chemotaxis is reducible to a non-homogeneous Markov process.

## Introduction

Decision making is de1ned as choosing a course of action from a set of possibilities (***Kitajima and Toyota, 2013***). Biological systems have to cope with both internal and external perturbations and make the “right” decisions amidst this pandemonium. The decision-making machinery is shaped by natural selection to 1t the conditions of its environment (***Tagkopoulos et al., 2008***; ***Mitchell et al., 2009***). Motility can be viewed as a decision-making process that bene1ts the living cell by enabling it to 1nd resources in its niche more efficiently (***Xie and Wu, 2014***). Cell motility requires sensors to monitor the environment, actuators to act upon the incoming information, and an network to process that information.

The majority of bacteria are motile, swimming being its most common form (***Jarrell and McBride, 2008***; ***Lauga, 2016***). Early studies revealed a substantial amount of variation in motility of clonal cells as they navigate a uniform environment (***Dufour et al., 2016***). In a homogeneous environment with uniformly-distributed resources, all decisions apropos of motility would be equally likely to be taken -i.e., *random walk*. Consequently in such circumstances, a motile cell would randomly navigate the environment. Having encountered a non-uniform distribution of resources in the environment, a motile cell will move to more resource-rich areas - i.e., the default *random walk* turns into *biased random walk*.

Chemotaxis, the ability of bacterial cells to sense chemical cues in their environment and move accordingly, predates the divergence of the eubacteria from the archaebacteria (***Woese and Fox, 1977***). The steps taken in chemotactic behavior, to seek attractants and avoid repellents, can be seen as a chain of biased random steps. To illustrate this point, we can focus on *Escherichia coli*. *E. coli* detects the concentration of chemoattractants in its vicinity via an array of sensors, processes the sensory data via a sensory network, and swims accordingly using its flagella (***Sourjik and Wingreen, 2012***; ***Frankel et al., 2014***). Following a trail of chemoattractants to get to their source is seemingly an insurmountable obstacle for *E. coli*, since their small size means that the difference between the amount of chemoattractants around its head and its tail would not be meaningful, and, consequently, useless in 1nding the correct direction. In reality, by rotating its flagella clockwise (CW) or counter clockwise (CCW), *E. coli* runs and tumbles through the environment (***Wadhams and Armitage, 2004***; ***Shimizu et al., 2010***).

Many mathematical models have been developed to understand the bacterial chemotaxis (reviewed in (***Tindall et al., 2008***)). The early models focused on the adaptive behavior of individual bacteria in different environmental conditions at a macroscopic level (***Segel, 1976***; ***Spudich and Koshland Jr, 1976***; ***Block et al., 1982***, ***1983***). Some models (e.g., ***Goldbeter and Koshland Jr (1982))***, used ordinary differential equations to describe the bacterial response to a gradient of chemical stimulants. Some used the Ising model (***Shi and Duke, 1998***; ***Duke and Bray, 1999***; ***Shi, 2000***, ***2001***, ***2002***; ***Guo and Levine, 1999***, ***2000***), others utilized an individual-based approach (***Frankel et al., 2014***; ***Niu et al., 2013***), and some emphasized the hydrodynamic aspects of swimming (***Elgeti et al., 2015***) and the role of drift versus diffusion (***Chatterjee et al., 2011***).

Most theoretical models of chemotaxis are limited to incorporating a single motor or simply assume that all cells have a single flagellum (***Bray et al., 2007***; ***Kalinin et al., 2009***; ***Matthäus et al., 2009***; ***Jiang et al., 2010***; ***Flores et al., 2012***; ***Kanehl and Ishikawa, 2014***). Constructing a comprehensive model of bacterial chemotaxis from the single-flagellum state has remained out of reach (***Mears et al., 2014***). What is more, most models do not explain how macroscopic chemotaxis behavior arises from the fundamental laws of chemistry and physics. In this paper, we propose a model that reduce chemotaxis to simple phenomena.

In our **S**tochastic **M**ulti-**L**ayer (SML) model, *E. coli* is treated like a minute biological submarine. This nano-submarine is propelled by an average of six flagella in low Reynolds’ number regime. Our model attempts to offer a comprehensive description of chemotactic behavior in *E. coli* by breaking this complex process into three levels. In the first level, chemoattractants react with the receptors, causing molecular events in the cell that can result in the rotation of each flagellum. The sensory network determines the direction and rotation rate of each flagellum. In the second level, each flagellum generates a force and the resultant force of all flagella causes the *E. coli* to move in the direction of this force. In the third level, the combination of different force vectors of each flagellum provides a range of direction and length of movement in each step - i.e., the behavior of the bacterium emerges from the chemical and the physical levels. At each step, as the concentration of chemoattractants sensed by the bacteria changes, so does the distribution of probability of all choices, i.e., the direction and the distance of travel at that step.

This hierarchical stochastic model is designed to model biochemical processes of chemotaxis within individual cells and the associated motion of cells within a 2*D* environment. This model can paint an *end-to-end* picture of chemotaxis and reveal its underpinning molecular mechanisms. In our view, while to details of our three-level model re2ects the intricacies of the biology, its behaviour would be indistinguishable from a **n**on-**h**omogeneous **M**arkovian **r**andom **w**alk (NHMRW) process.

## Results

### The Macroscopic Behavior of the SML model is indistinguishable from a NHMRW

We characterize the macroscopic behavior of the SML model by comparing it with a random walk-as an unbiased foraging process-and the non-homogeneous random walk. The qualitative behavioral difference with the random walk is clear (Figure 1).

**Figure 1.**
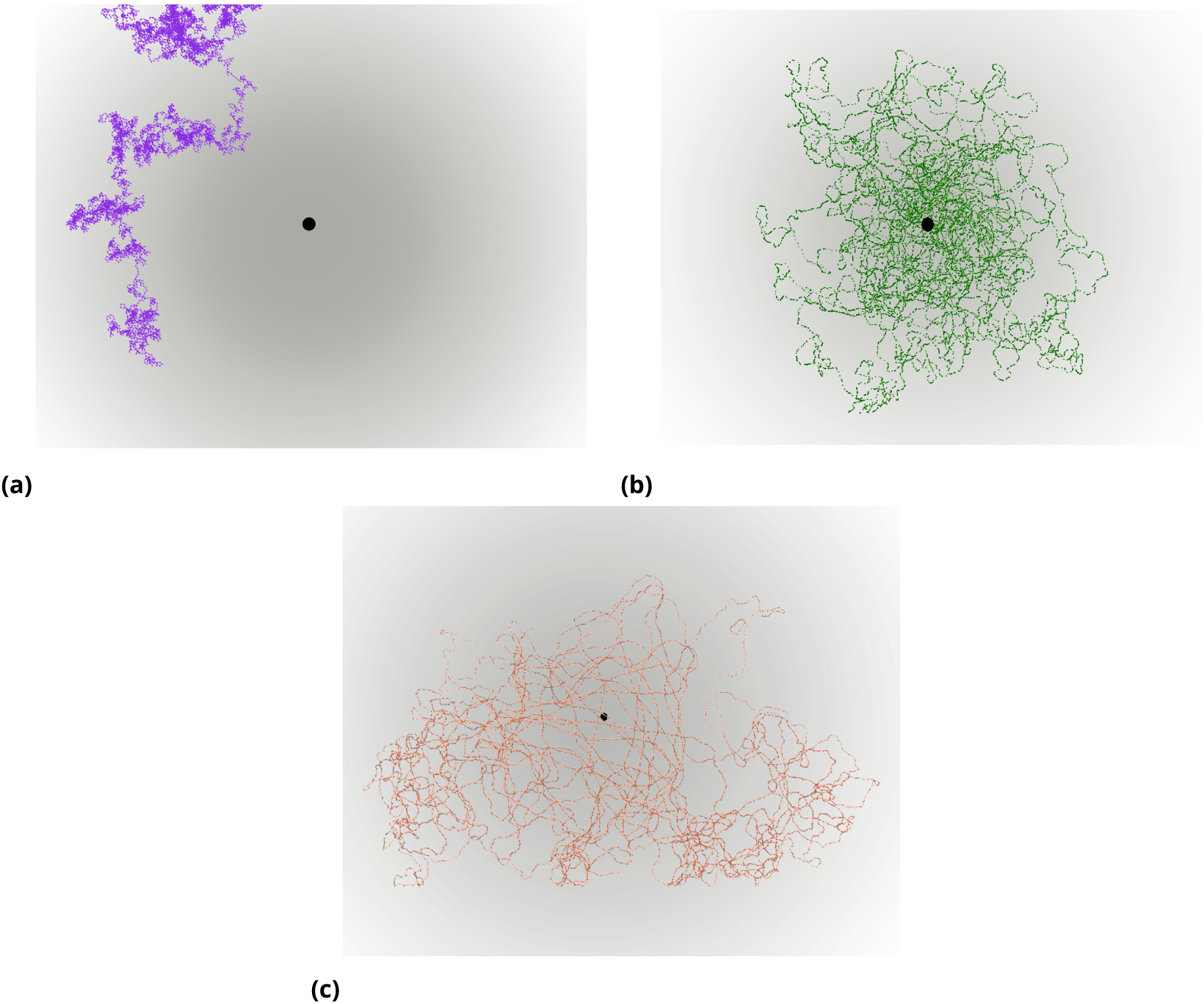
Trajectories of the random walk (left), the SML (right), and the NHMRW (buttom) models in a two-dimensional space. Results based on 1000 trajectories for each models.

While cells in the random walk process are roaming around and spent most of their time in a random location without any correlation between the concentration of the nutrients and their location, the bacterium in the SML model revolves around the high concentration area, i.e., nutrient concentration values and spatial movement direction are strongly correlated.

However, there is a behavioral similarity between the SML and non-homogeneous random walk model (*NHMRW*): in the *NHMRW*, similar to the SML model, the movement to areas of higher chemoattractant concentration is more preferable (Figure 1). Both models show the same dynamical behavior for *ϕ* as the concentration of nutrients varies. Moreover, the mean deviation angle, namely, 64°, is consistent with the 1ndings of the experimental observations by ***Turner et al.*** (***2016***) (table 2). The experimental mean value for *E. coli* tumble angle was found to be around 68° (***Berg et al., 1972***), and 64° (***Turner et al., 2016***).

**Table 1.** Averaged number of simulation steps for particles to reach the maximum concentration.

| value | SML | NHMRW | Random Walk |
| --- | --- | --- | --- |
| mean | 50.5 | 48.3 | 422.1 |
| std | 29.01 | 30.7.5 | 387.1 |

**Table 2.** The average change in the direction of particles in each simulation step in comparison with the experimental data sets from ***Berg et al. (1972)*** and ***Turner et al. (2016)***, respectively.

| value | SML | NHMRW | Random Walk | data set |
| --- | --- | --- | --- | --- |
| $\varphi$ | 57.5° | 59.9° | 38.9° | (68°, 64°) |

Bacteria tumble more frequently as they move in areas with nutritional de1ciency, to increase the chance of survival (Figure 4). Qualitatively, tumbling frequency in the SML model agrees well with the experimental reported data sets (***Balázsi et al., 2011***; ***Mittal et al., 2003***). In this work, this quantity is calculated by scaling the concentration to the range [0, 10] and considering the steps with deviation angle greater than 25° as tumble.

### The rotational directions of all flagella combined determine the direction of movement

In *E. coli*, the flagellar rotation is the driving force behind motility. In response to the changing nutrient concentration, the chemical network in the cell regulates the rotational direction of each flagellum (*CW*⇄*CCW* rate). Consequently, the cell is capable of adjusting its mean speed and the distribution of tumbling angles to position itself more effectively. The switching rates of flagellar motors from the SML model are comparable with reported values in ***Mears et al.*** (***2014***) (table 3).

**Table 3.** CW and CCW switching rates as comparison with the experimental outputs in (***Mears et al., 2014***)

|  | SML | experiments |
| --- | --- | --- |
| $CW \Rightarrow CCW$ | 0.21 % | $0.26s^{-1}$ |
| $CCW \Rightarrow CW$ | 0.79 % | $1.7s^{-1}$ |
| Average CW bias of motors | 0.21 % | 0.13 |

### The chemical network results in directional sensing

The concentration of CheY-p plays a major role in the signaling network of *E.coli* chemotaxis. A well-known feature of the chemotactic network is the sigmoidal relation between the direction of the flagellar rotation and the CheY-p concentration (***Yuan and Berg, 2013***). A comparison between the CheY-p concentration from our Gillespie simulation (the chemical level) and the experimental data in ***Mears et al.*** (***2014***); ***Lele et al. (2015)***; ***Terasawa et al.*** (***2011***) is given in table 4. Any change in the CheY-p concentration would cause a change in the probability of rotational direction of flagellar motors (***Sagawa et al., 2014***). On average, 13 ± 7 and 2 ± 4 CheY-p molecules bind to a flagellar motor during CW and CCW rotation respectively (***Fukuoka et al., 2014***; ***Segall et al., 1985***).

**Table 4.** Comparison between CheY-p concentration calculated by SML model and experiment (***Mears et al., 2014***).

|  | SML | experiments |
| --- | --- | --- |
| Mean([CheY-p]) $\mu M$ | 1.69 | 2.59 |
| Var([CheY-p]) $\mu M^2$ | 0.76 | 1 |

## Discussion

Gazing upon the movement of living entities invariably instigates a chain of thorny questions regarding the nature of movements. As Aristotle observed, in his *De Motu Animalium*, “it remains to inquire how the soul moves the body, and what is the origin of movement in a living creature” (***Barnes (1995)***, p.2383). It is tempting to scoff at the idea of an *èlan locomotif* pushing a living entity forward, but one can hardly fault an observer studying the movement of a bacterium under the light microscope for inferring a certain intentionality from the movements of that organism.

The movement of a bacterium, such as *E. coli*, can be characterized as a series of “decision”. Throughout this work, we have used decision making as a mere shorthand to denote change in the behavior of the organism caused by processes at the molecular level; a kind of decision making that is devoid of any intentionality and comprehension. To achieve this, our SML model simulates the CW and CCW rotations of flagella as a function of the concentration of chemoattractants in the environment. The comparison between the SML model and a random walk alternative vividly demonstrates the efficacy of our model, whereby our digital *E. coli* spends signi1cantly more time at zones with higher chemoattractant concentrations (fig. 2 and table 1). The SML model is unique in that it simulate a bacterium with six functional flagella. This level of realism in simulating cell motility is absent from similar studies, which are content with including a single fellaglla. This melange of modest, yet unprecedented, cellular realism, and a stochastic approach to simulating molecular process, a salient feature of the SML model, is an attempt to re2ect the inherent complexity, as well as the innate stochasticity, of living entities.

**Figure 2.**
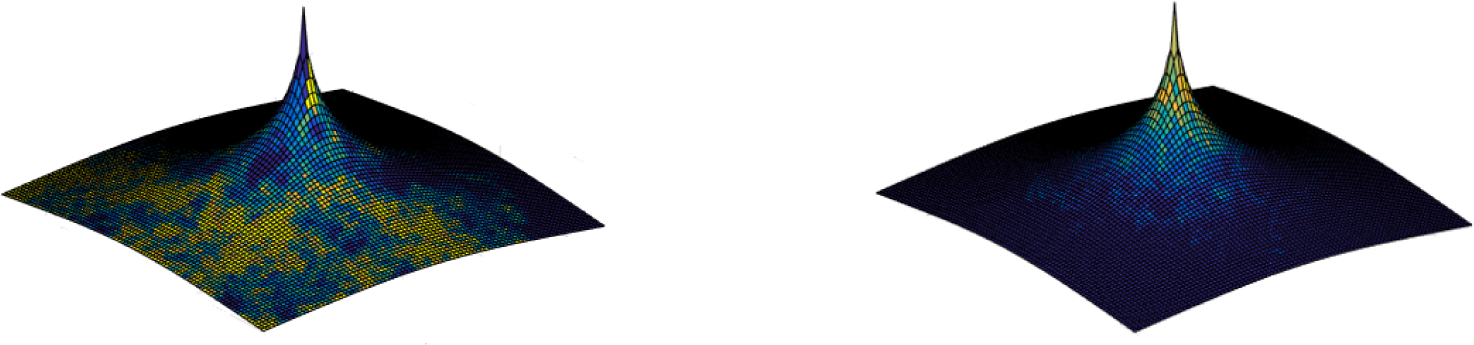
Trajectories of 1000 simulations in the random walk (left) and the SML model (right). Starting points are randomly chosen point, with random angles, 300 units away from the nutrient source. Each run will end if the particle 1nds the nutrient resource or the number of simulation steps exceeds 3000. The height re2ects the density of particles.

The probabilistic nature of the SML model means that chemoattractant concentrations below the sensitivity of this model results in a behavior indistinguishable from the random-walk alternative, but as the chemoattractant concentration increases, so does the bias of *E. coli* movement (fig. 3). The comparison between the SML model, the Markovian model, and the experimental data indicates the similarity between the SML model and the way *E. coli* behaves in real life (table 2). The mean angle of movement in our model is slightly different from experimental data (57.5° versus 64° (***Berg et al., 1972***) and 68° (***Turner et al., 2016***)). This slight discrepancy can be partly attributed to neglecting near-zero angles in the experiments; a similar discrepancy can be observed between Turner et al.,2016 and Berg et al.,1972, where different thresholds to distinguish running from tumbling were used. In this work, tumbling, de1ned as a movements with angle *>* 25°, is determined by the resultant movement vector of all flagella. The movement vector for each flagellum depends on its direction of rotation, itself the function of the number of Chey-P proteins attached to it. Treating the movement of each flagellum independently is in accordance with studies such as Mears et al.,2014.

**Figure 3.**
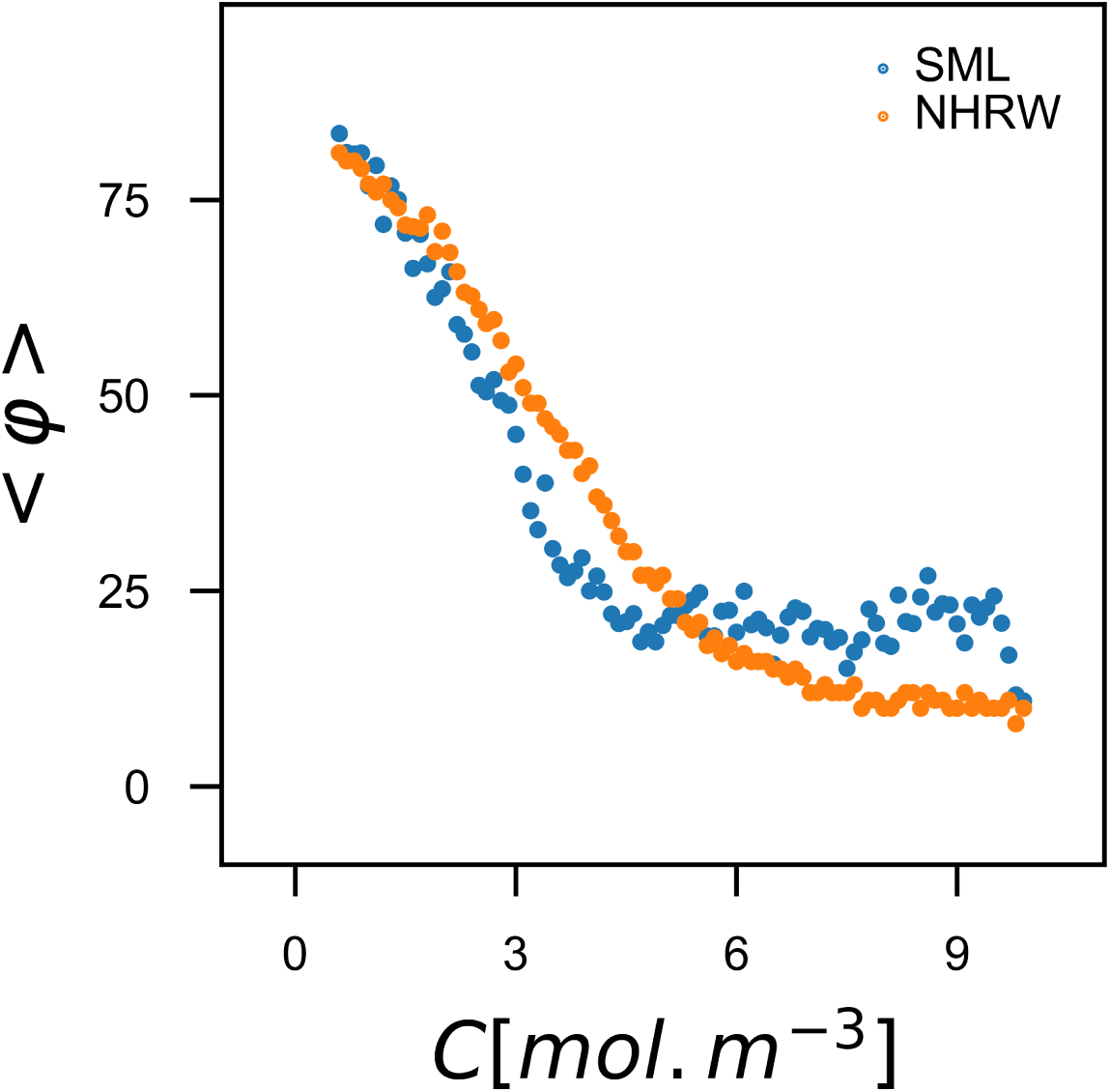
Changes in the movement direction of the particles at each simulated step with respect to their previous direction. As glucose concentration increases, *E. coli* frisks less and waggles more, i.e., there are less variations in the direction of movement. The results are averaged over 1000 simulations.

**Figure 4.**
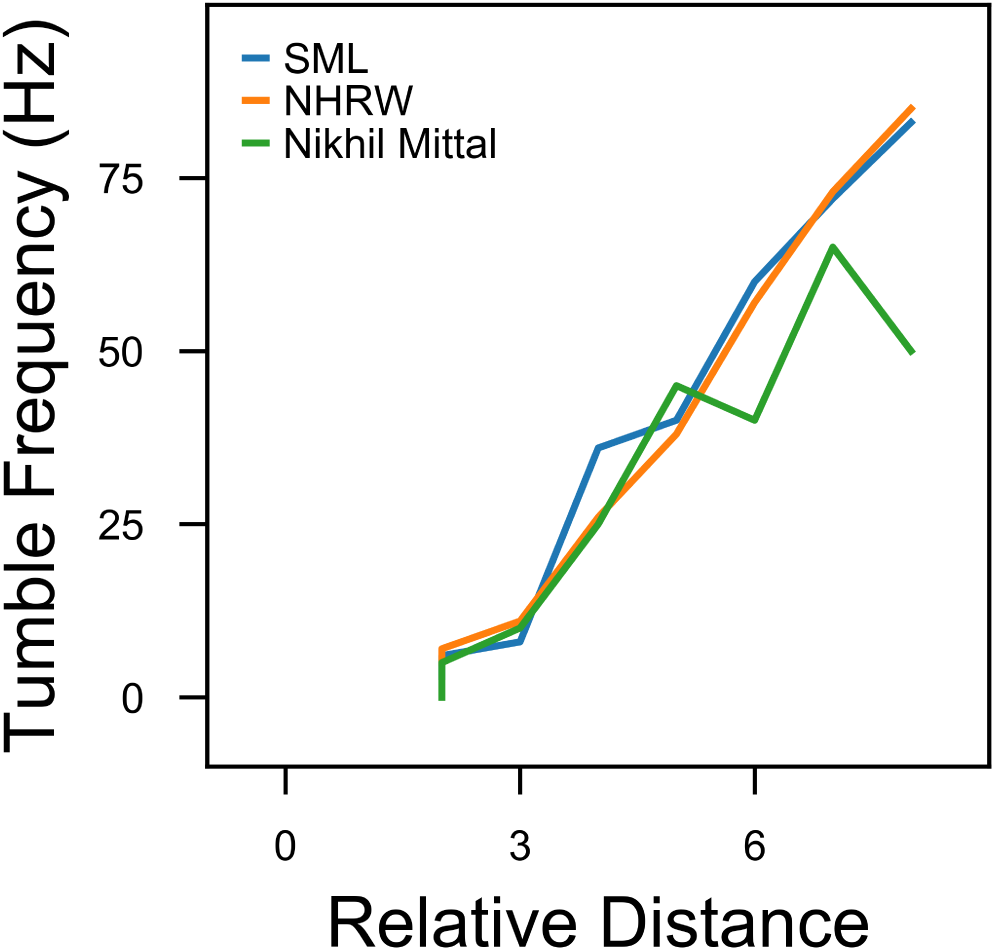
According to the results in ***Mittal et al.*** (***2003***), the tumble frequency positively correlates with the distances from the resource. The same pattern is observed in the SML model, where the chemoattractant concentration decreases radially away from the resource, hence, *E.coli* would change its direction more frequently. Following ***Mittal et al.*** (***2003***), we considered angles greater than 25° as tumbling.

**Figure 5.**
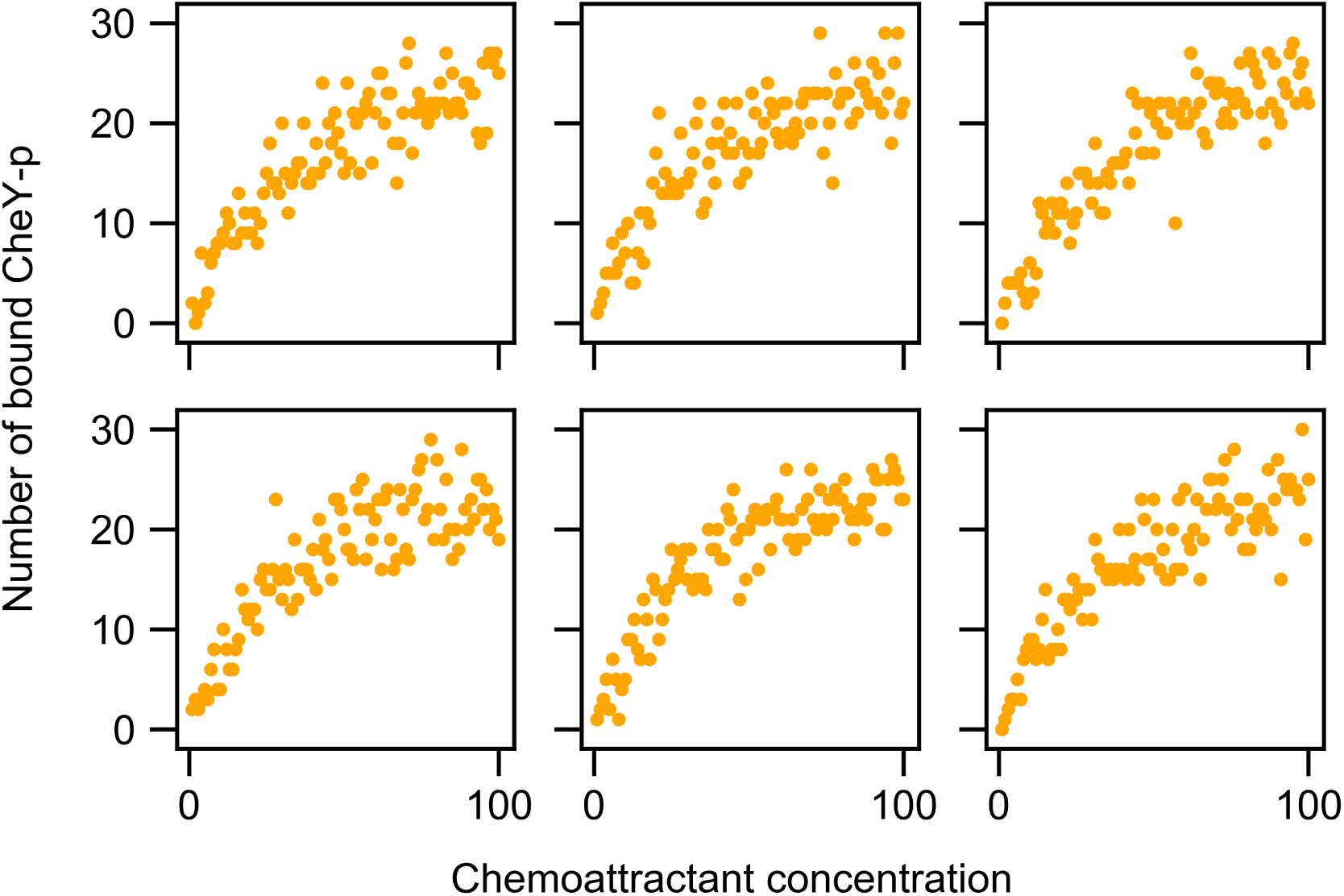
The number of *CheY* molecules bound to each flagellum in different chemoattractant concentrations. The binding of the phosphorylated CheY (CheY-p) to flagellar motors increases the probability of a transition from CCW to CW rotation of the motor.

The similarity between the SML model, with all its molecular accoutrements, and the Markovian model, might seem quite irrelevant on the surface: The Markov process simply captures the macroscopic behavior, while utterly oblivious to the intricacies at the cellular level. However, this similarity can be interpreted in a starkly different manner: our Markovian model, though deeply devoid of any biological realism, can keep up with the SML model, in describing the macroscopic level. While no molecular machinery can be gleaned from the Markovian model, it does enable us to predict the behavior of a living entity in an accurate fashion.

Why should we bother with a hierarchical model, combining physical and chemical levels to investigate the movement of a cell? This layered approach allows for different experimental measurements to be incorporated in a singular model. In addition, by utilizing an approach similar to the SML model, one can compare results studies at the chemical level (e.g., Shimizu et al,2010 and Sourjik and Berg, 2002b) with treatments of the physical level (e.g., Rodenborn et al.,2013). A detailed model of any cellular behavior, such as cell movement, enables us to make testable predictions as well.

Our attempt here was to reconstruct the behavior of a complex entity, namely a free-living prokaryote, by incorporating a detailed chemical level to a physical level. It is only natural that the exact behavior of the SML model would change if you change the parameters (for example, the rate of Chey-P attachment to a flagellum reported by ***Hosu and Berg*** (***2018***) - which we used in our model - differs from that of ***Bai et al. (2012))***. More generally, modelling is a process of simpli1cation and these simpli1cations, such as treating *E. coli* cell as a sphere, might result in further deviations from reality.

Despite all the possible 1ne tunings and the inescapable reductionism inherent in a model of movement, we can qualitatively answer what Aristotle asked about the seemingly magical feature of living bodies, i.e., to move without being moved. It is not a “soul” that moves the entity forward, but the stochastic chemical reactions that become biased enough in response to the environmental cues. Here, we have shown that a seemingly complex feature of *E. coli*, namely its ability to explore the environment, simulated here using a multi-layered model, can be easily reduced to non-homogenous Markov process. A biased random walk that at the macroscopic level that is so deceptively directed as to imply a sort of intent to the untrained observer. But the reality is far more pedestrian, and yet far more majestic. The seething chemical soup within a living cell can result in a behavior -i.e., movement-that seems utterly alive.

## Methods

To simplify the implementation of the cell migration and mobility, we mainly focus on the cell chemotaxis without considering cell division; moreover, we consider *E. coli* as a sphere with non-interacting flagella.

### Chemical Intracellular Interactions

The eukaryotic means of detecting the chemical gradient in the environment directly is not useful to bacteria given their comparatively diminutive size. In fact, many chemotactic bacteria navigate by measuring temporal changes in concentration as they swim. he classical stochastic simulation algorithm (SSA) by Gillespie and its modified versions are widely used to simulate the stochastic dynamics of biochemical reaction systems. It has, however, remained a challenge to implement accurate and efficient simulation algorithms for general reaction schemes in growing cells (***Yu et al., 2010***).

Figure 6 shows the schematic view of *E. coli* and its chemotactic chemical network. *E. coli* can merely sense the environment by an array of chemoreceptors to perceive the concentration of chemoattractants. This sensory array triggers the inner network in *E. Coli*. The probability of CheY-p protein binding to motors is regulated by the input of inner network as the Navigation system. We apply a radially-decreasing nutrient gradient away from a local resource, in which the attractant concentration,

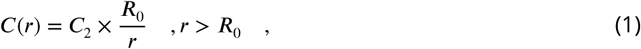

is constant within a ball of radius *R*_0_ = 100*µm*.

**Figure 6.**
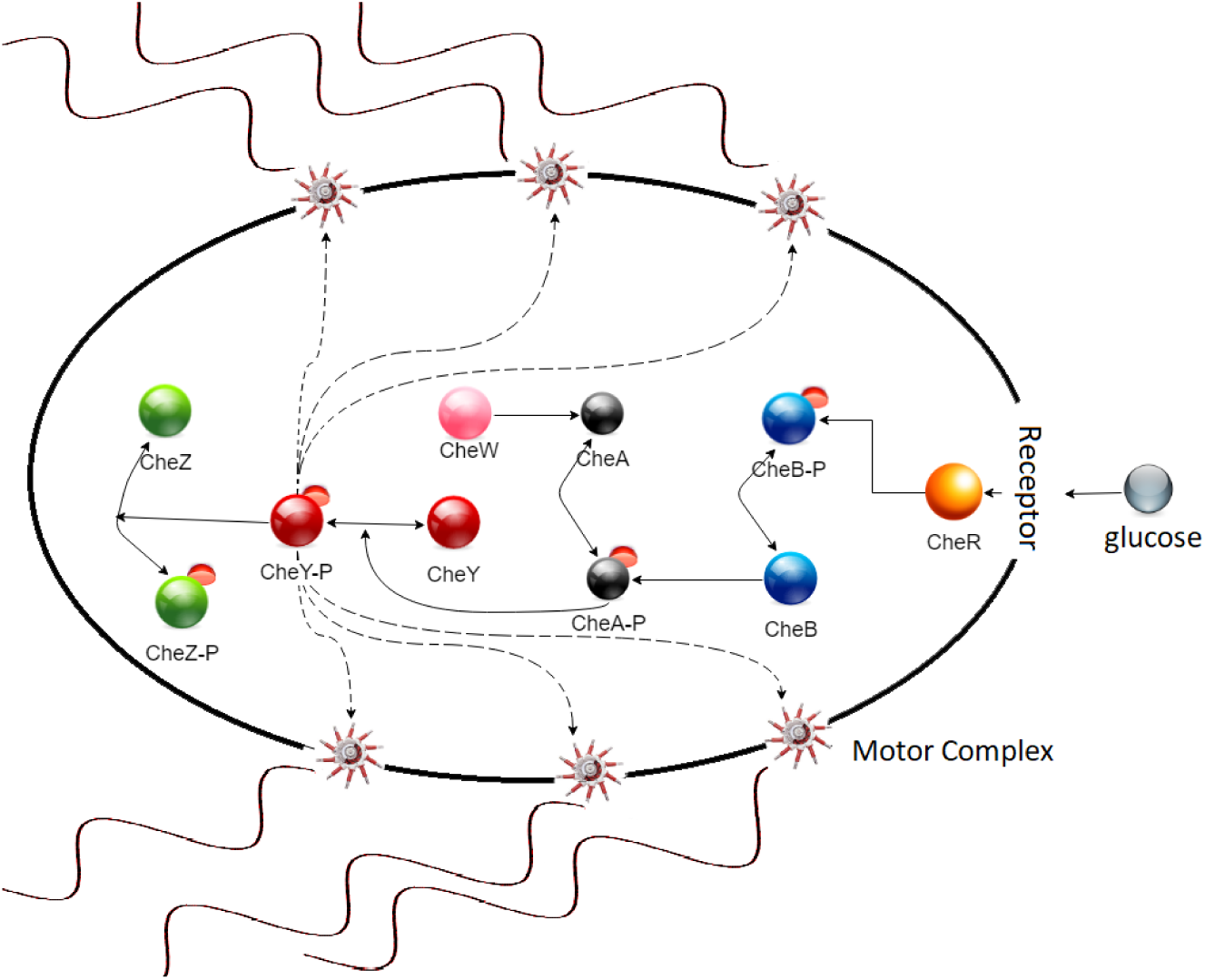
Schematic diagram depicting the body of *E. coli* and the chemical network inside the cell. Receptors sense the chemoattractant concentration from the environment, then trigger signaling cascade in the chemotactic network. The number of CheY-p proteins bound to the motor complex of each flagellum– resulted from internal chemotactic network– determines the rotational direction of that flagellum.

**Figure 7.**
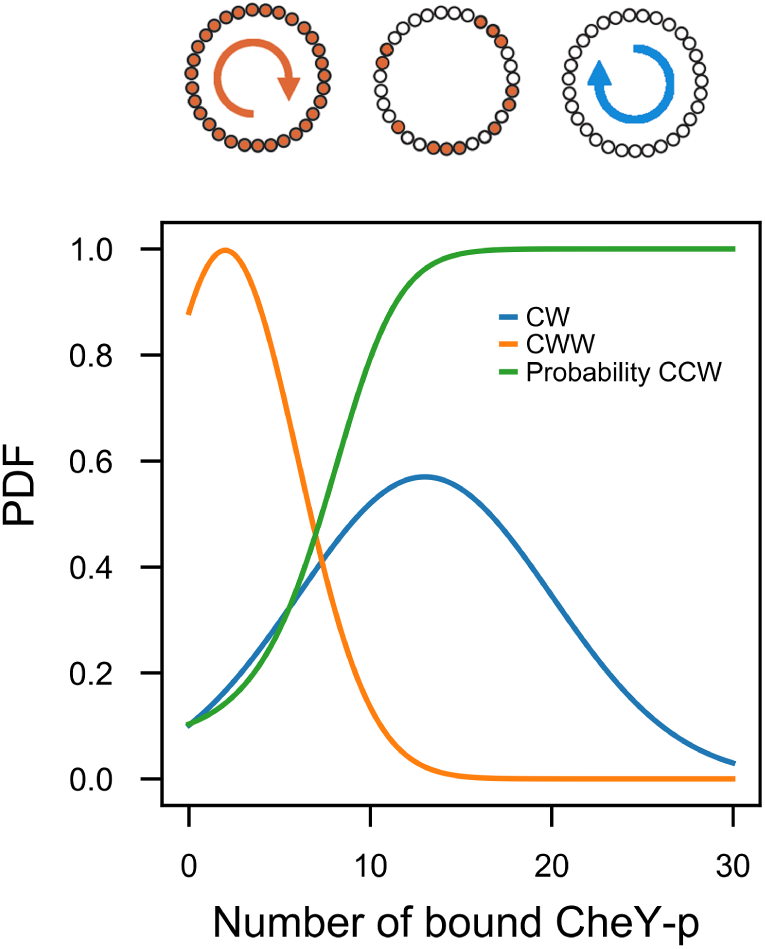
The binding of CheY-p to the flagella motors (depicted as filled in circles) increases the probability of a transition from CCW to CW rotation of the motor. Correlation between CCW/CW rotation of flagellum and the number of bound CheY-p. The binding of CheY-p to the flagella motors increases the probability of a transition from CCW to CW rotation of the motor.

As a cell swims, chemoattractant molecules bind to the receptors on the cell surface; therefore, a signal from receptors would be transmitted, stochastically, through a biochemical network to one or more of the flagellar motors, which controls the speed and direction of the flagellar rotations. We assume the following dependency for the sensitivity of these receptors to the chemoattractant concentration, *c*,

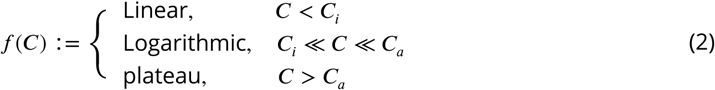

in this work, the values for the threshold concentration, *c*_*i*_ = 0.0182*mM* and *c*_*a*_ = 3*mM*, are assumed according to ***Shimizu et al. (2010)***. Thus, as a cell approaches a high concentration of nutrients, its sensitivity decreases, which effectively increases the value of *CheR*, see chemical network in figure 6. In order to fulfill the logarithmic dependency,

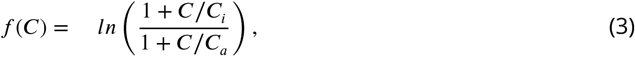

a localized attractant resource was assumed.

Chemoreceptors in *E. coli* are coupled to the flagella by a phosphorylated intermediate, CheY-p. CheY-p activity can be inferred from the rotational bias of the flagellar motors, although the motor response is stochastic and limited to a narrow physiological range (***Sourjik and Berg, 2002***).

### Chemotactic Network

The signal transduction between receptors and flagellar motors is controlled by a set of well-de1ned intracellular protein-protein interactions (***Wadhams and Armitage, 2004***). The core of the network is a two-component signal transduction system that carries the chemical information, gathered by transmembrane receptors, to flagellar motors responsible for the cell propulsion (Figure 6). A second group of proteins allows cells to physiologically adapt to the changing levels of the background signal, enabling them to track signal gradients over many orders of magnitude.

While different receptors allow cells to sense different signals, all signals are then processed through the same set of cytoplasmic proteins, responsible for signal transduction and adaptation. Signals can vary in time, space, and identity; consequently, this horizontal integration may impose incompatible demands on the regulation of these core *decision-making* components. In this study, we applied a simplified abstract network in which a number of intracellular proteins– known as chemotaxis (Che) proteins– provide the necessary signaling cascade which links the membrane receptors to the flagellar motors (Figure 6). CheW and CheA are chemotactic proteins bound to the receptors. CheW is thought to act as a link between the receptors and CheA. In addition, CheA appears to directly interact with the receptors. To bring about tumbling, the receptors activate CheA autophosphorylation on a conserved histidine in response to decreased attractant or increased repellent concentration. One of the phosphoryl groups is transferred to CheY. CheY-p shows a reduced affinity for CheA and a higher affinity for the flagellar motor protein FliM. Therefore, it diffuses through the cytoplasm to the motors. CheZ acts to dephosphorylate CheY-p at the receptors to regulate the rate of signal termination (***Tindall et al., 2008***).

#### Flagellar Rotation

The rotation of the bacterial flagellar motors is controlled by the above mentioned signal transduction pathway. The switch from CCW to CW rotation is triggered by binding of the signaling protein CheY-p to the motor. The direction of flagellar rotation and the amount of the generated torque is regulated by a complex at the bottom of the basal body called the switch complex, constructed from FliG, FliM, and FliN proteins (***Sarkar et al., 2010***). The distribution of CW and CCW switching intervals depends on parameter sets for the volume of localization and the number of localized molecules (CheZ) (***Yu et al., 2010***). A discrete stochastic model captures the 2uctuations in the rotational direction of the flagellar motors (CW ⇆ CCW), in which the random binding of CheY-p proteins to the motor-binding protein FliM leads to the motor switching from either a CCW rotation to CW or vice-versa. The probability of CCW rotation is assumed proportional to the concentration of CheY proteins,

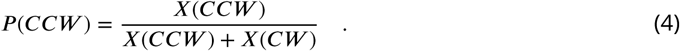

The parameters in this model have been chosen based on the available experimental data (***Fukuoka et al., 2014***). On average, 13 ± 7 and 2 ± 4 CheY-p molecules bind to a flagellar motor during CW and CCW rotation, respectively,

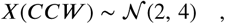

and

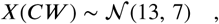

where 𝒩 (*i, j*) means *X* has normal distribution with *mean i* and *variance j*. The number of CheY-p molecules binding to CW rotatory motor plateaus at about 13 molecules, instead of saturating at 34 molecules which is the number of FliM subunits (***Hosu and Berg, 2018***).

Based on our assumptions, there is no correlation between the rotational direction of flagellar motors. Any interaction between flagellar motors and CheY-p– generated by chemoreceptors– would be inhibited by the CheY, which is distributed uniformly throughout the cell. Rotational direction of motors is dictated by CheY-p and the motor closer to the chemoreceptor patch would switch earlier than a motor farther from it (***Terasawa et al., 2011***).

### Physical Movement Mechanism

The cell body is considered as a solid sphere of diameter *R*_*b*_ = 9Å with six flagella of length *L* = 6.6*pm* randomly distributed on the cell membrane, see figure 6. Flagellar diameter depends on its state (***Rodenborn et al., 2013***) (table 5).

**Table 5.** Parameters used in Eq. 5

| Parameter | Value |
| --- | --- |
| R(CCW) | $0.195 \pm 0.25 \mu m$ |
| R(CW) | $0.210 \pm 0.25 \mu m$ |
| $\lambda/R$ | 11 |
| $L/\lambda$ (CCW) | 2.8 |
| $L/\lambda$ (CW) | 2.7 |
| $\tan(\alpha)$ | $2\pi R/\lambda$ |

Flagellar motors are capable of rotating with angular speeds of up to hundreds Hertz, enabling the bacterium to propel itself through the extracellular environment. To model the role of the flagella in the motility of *E.coli*, it requires to calculate the total force generated by all flagella. At each time step, each flagellum exerts a force with magnitude and direction driven by its rotatory motor which is the result of the stochastic chemotactic network. Consequently, the total force– applied on the cell body by all the flagella– would be a stochastic quantity.

The helical flagella are driven by a rotary motor embedded in the wall of the body, spinning with angular speed Ω_*m*_ relative to the body (***Hu et al., 2015***). Each *flagellum* by its rotation causes a drag force *F*_*i*_, *i* ∈ [1 : 6]. For simplicity’s sake, we assume flagella are distributed around the body in only two directions relative to the cell surface: 1) when a flagellum rotates CCW, it would be aligned parallel to the cell body, as well, its generated force; 2) when it rotates CW, the force is perpendicular to the cell body. At low Reynolds numbers (the linear Stokes equations), by applying an external force *F*_*i*_, a solid body will move with velocity *U*, which is proportional to the angular speed (***Lauga and Powers, 2009***; ***Lauga, 2016***), by assuming *L* » *R*_*b*_,

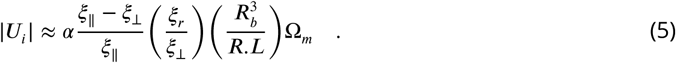

*ξ*_‖_ and *ξ*_┴_ are drag coeZcients on the directions parallel and perpendicular to the flagellum, respectively (typically *ξ*_‖_/*ξ*_┴_ ~ 2). Movement direction of a cell, subject to applied forces by 6 flagella, is determined by the resultant velocity vector, 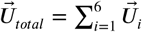.

Therefore, it would be possible to evaluate the direction (*φ*) and step length (*l*) of the next foraging step. *φ* is in line with the direction of *U*_*total*_ and by assuming a 1xed speed, *l* = *U*_*total*_ × *τ*_*step*_ (***de Lima Bernardo and Moraes, 2011***). According to the experimental results, run and tumble duration times are exponentially distributed with mean values 〈*τ*_*run*_〉 ~ 1*s* and 〈*τ*_*tumble*_ ~ 0.1*s*, respectively, (***Alon et al., 1998***). In our model we have only one type of movement, thus, for each time interval, we select a random number from the exponential distribution with 〈*τ*_*step*_ ~ 1*s*. By calculating *φ* and *l*, the next spatial position is taken as follow,

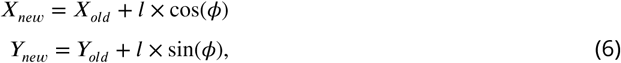

and the simulation will be continued the same way.

## Supporting Information

### Non-homogeneous Random Walk

In an abstract view, we modeled the chemotaxis of *E. coli* as a non-homogeneous random walk. In this stochastic process, in each simulation step, the probability distribution of speed, direction, and time interval for the next step are independent of each other. The time interval of each step (or waiting time until the next turning point) comes from an exponential distribution with mean value 〈*τ*_*step*_〉 1 ~ *s* and a normal distribution governs the speed of each step. In order to derive the direction of each step with respect to the previous step, *ϕ*, which is strongly in2uenced by the concentration of nutrients, we applied a Beta distribution function,

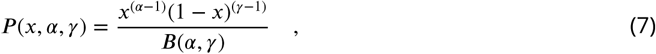

where *x* depends on the chemoattractant concentrations limited to 0 ≤ *x* ≤ 1. *B*(*α, γ*), Beta function (***Forbes et al., 2010***), is the normalization constant,

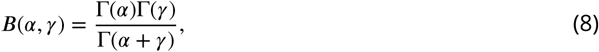

with *α, γ* > 0. In this model, we limited *ϕ* to [*ϕ*_*min*_, *ϕ*_*max*_]; the selected values for the upper and lower limits are according to experimental measurements (***Masson et al., 2012***). To simplify our calculation, we rescaled the limits of *ϕ* to [0, 1] range,

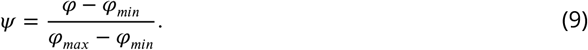

To specify the Beta function in Eq. 7, we need to assign *α* and *γ* values. To achieve the desired configuration for the distribution function of angels (***Masson et al., 2012***), we exerted identical values for *α* and *γ*, dependent on the chemoattractant concentrations,

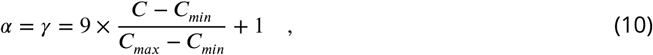

which are in the range of 1 to 10 (Figure 8) illustrates the Beta distribution for different values of *α* and *γ*. Based on Eq. 10, for the minimum values of the chemoattractant concentration, both *α* and *γ* are equal to 1 and the Beta distribution would be equivalent to the uniform distribution; hence, all valid angels in the range of [*ϕ*_*min*_, *ϕ*_*max*_] would have the same chance to be selected.

Whenever *E. Coli* receives the chemoattractant at the highest value, *α* and *γ* take the value of 10 and movements along an straight line are more probable. The receptors sense the intensity of glucose concentration by a Monod function (Eq. 3). The inner network determines the rotational direction of each flagellum by calculating the number of CheY-p proteins bound to the motors. In this model, the nutrient concentration, *C*, highly affects values of *α* and *γ*. Consequently, any variation in *α* and *γ* will change the probability distribution in Eq. 7. In fact, in the non-homogeneous random walk model, the decision-making network is modeled through probability distribution functions. Indeed, to perform the simulation, after generating a random number *x* from *p* (*x*) in Eq. 7, direction of movement, *ϕ*, could be derived as follow,

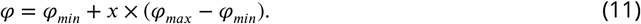

**Figure 8.**
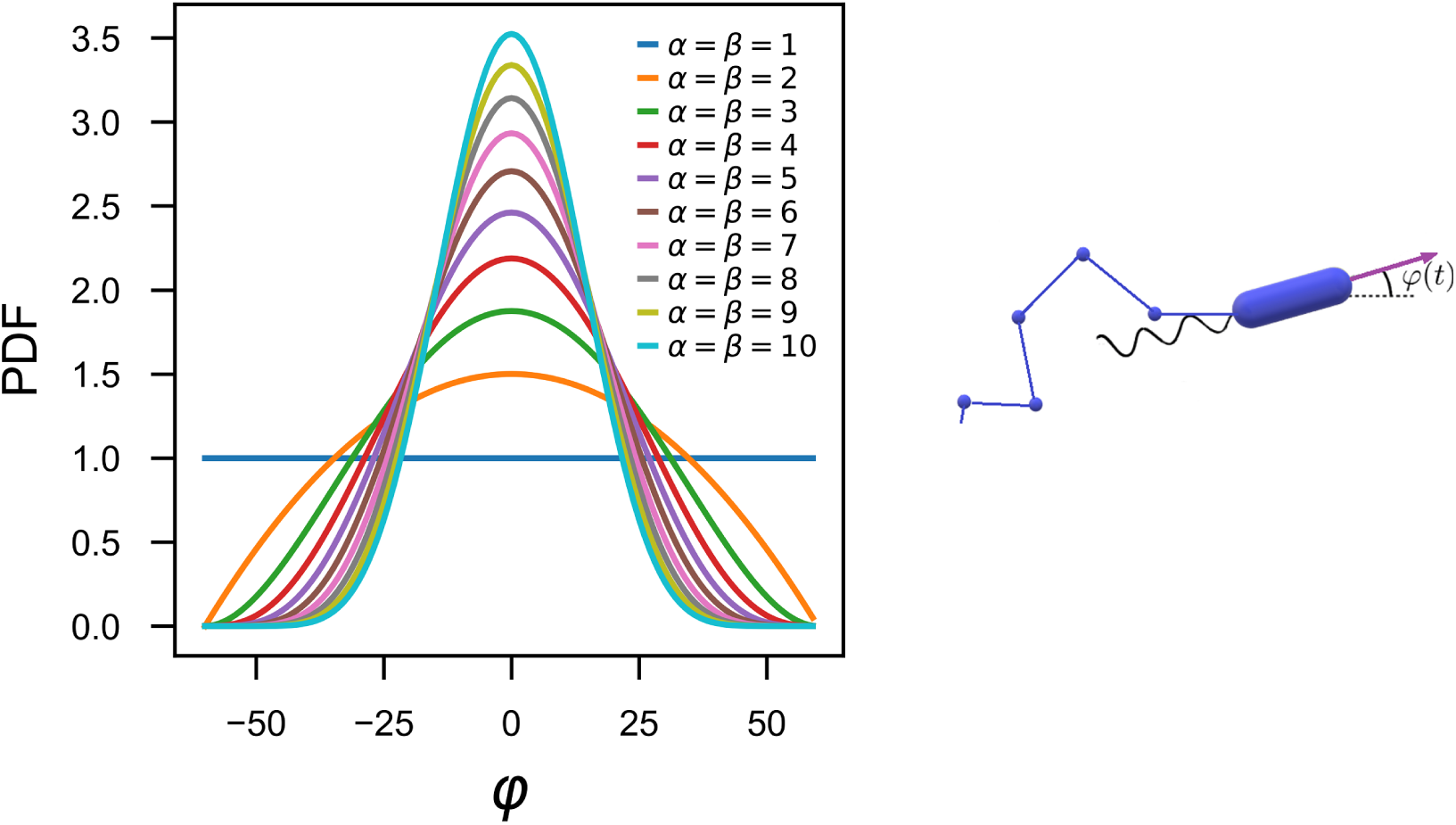
The concentration of chemoattractant strongly affect the distribution of the relative angle between two consecutive steps *ϕ*(*t*). Different values from the Beta distribution result different directions of movement for the cell (Right). The shapes of Beta distribution for different input sets, *α* and *β*, shown by *B*_*i*_ (Left). In an environment with uniform distribution of nutrients, the distribution of *ϕ* is uniform as well, the blue curve; i.e., every direction has the same chance of being chosen. As a result of a non-uniform distribution of nutrients, a normal distribution will emerge (the green curve).

**Figure 9.**
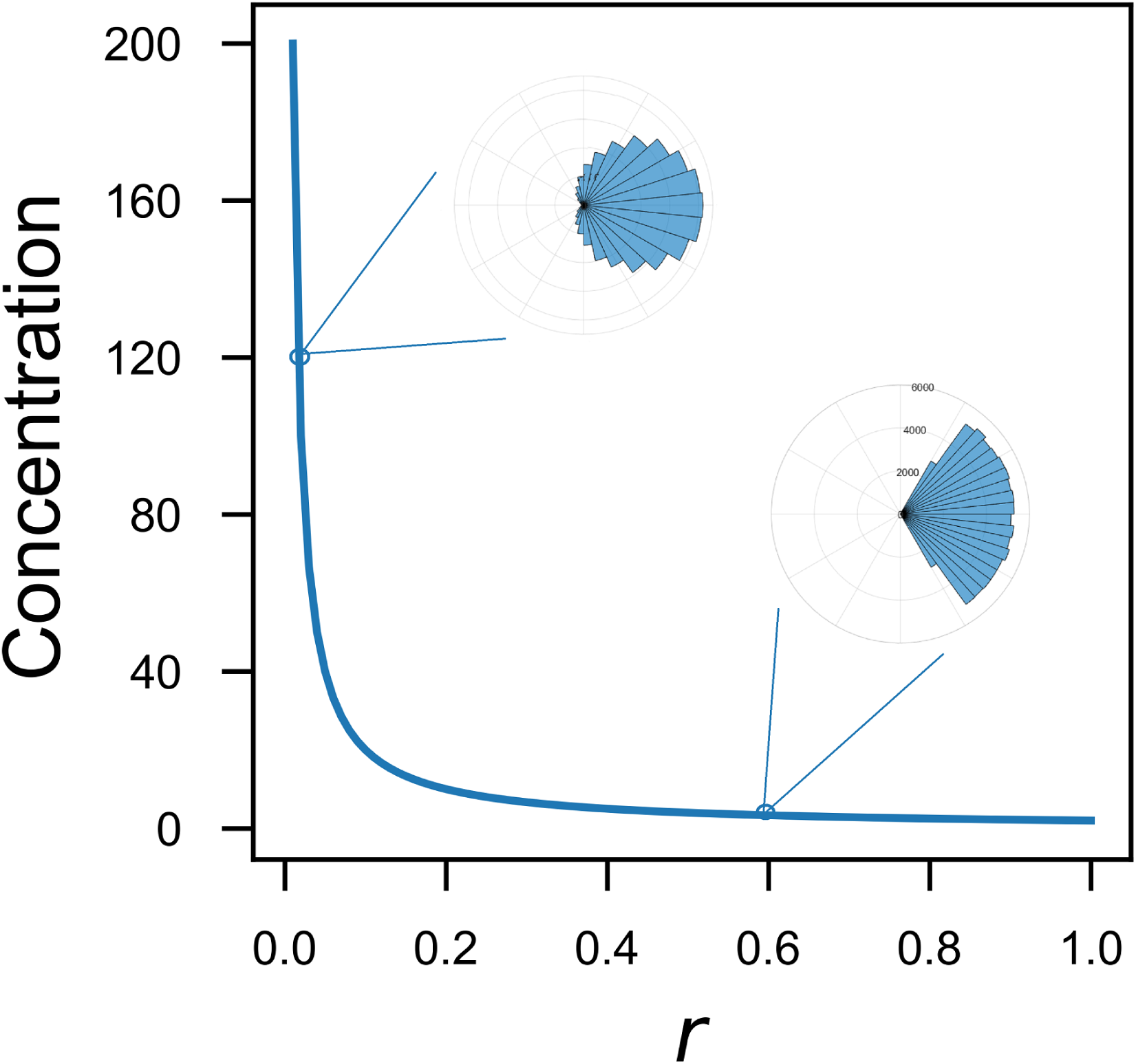
The concentration of the perceived nutrients by the chemotactic network (*r*) affects the distribution of angles. As the perceived concentration increases, so does the probability of smaller angels, i.e., movement closer to an straight line. However, there is not a straightforward linear relation between them since many parameters, including the sensitivity of the receptor (Eq. 3) play roles.

**Figure 10.**
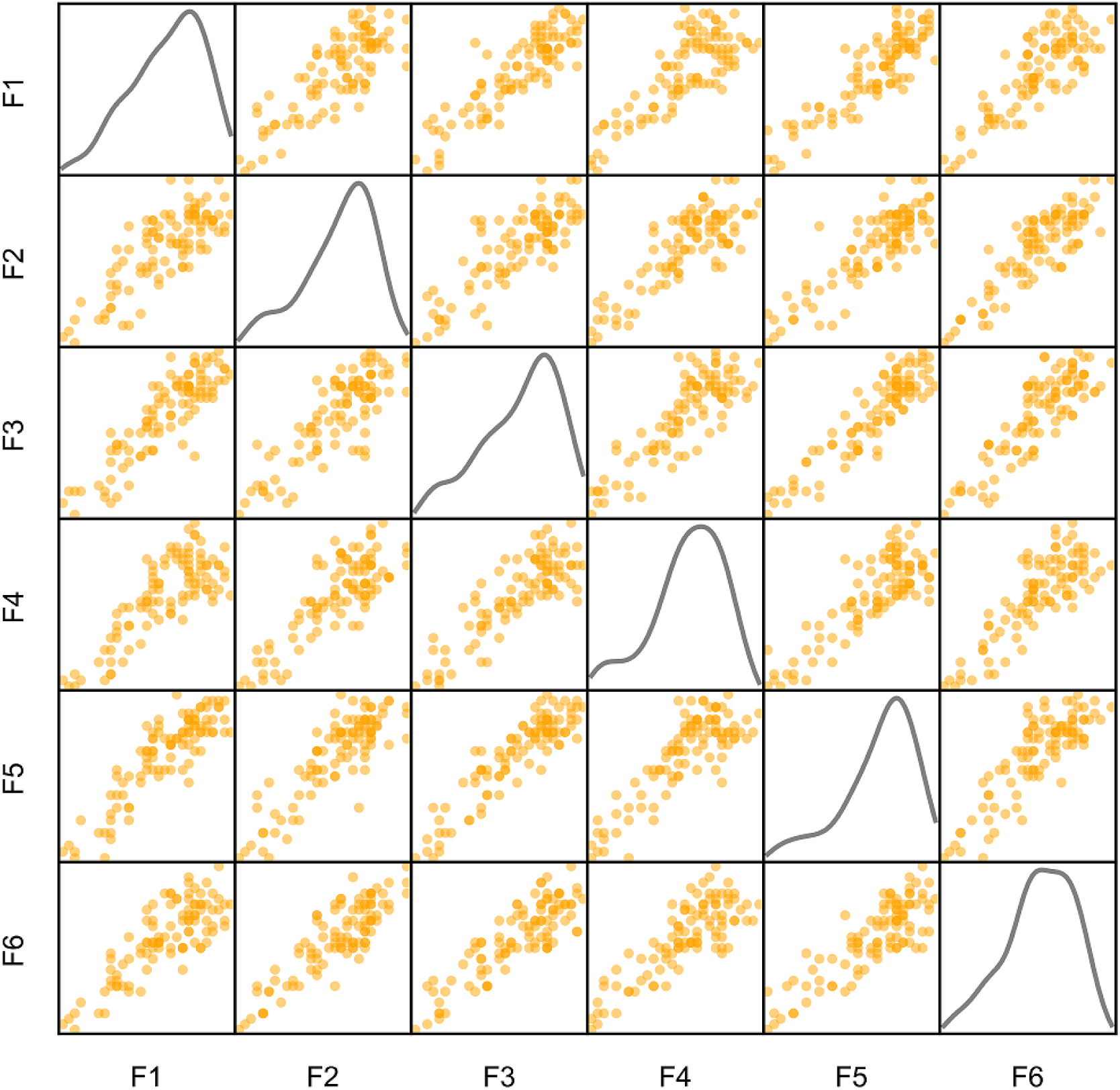
There is no absolute correlation between the flagella or the position of the flagella in the SML model. The diagonal figures are histogram plots of number of each flagella. Each figure in the *r* the row and *c* the column of the matrix shows correlation between flagella *r* and *c*.

A normal distribution 𝒩 (*µ*, 10) with mean *µ*, from Eq. 12, gives the speed of the next step.

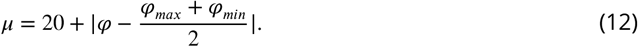

Knowing the speed, direction and duration of each step, we readily compute the next spatial position by Eq. 6.

*lagella-CheY-P* concentration changing dynamics

## Acknowledgments

We would like to thank Dr. Yazdan Asgari for his early works relevant to this study.

## Additional Information

### Author contribution

MS conceived the model and helped draft the manuscript. KK contributed to the initial idea. HS contributed to the physical layer of the model. SV constructed the model, performed the simulations, and drafted the manuscript. SV and AK visualized the data. MS and AK contributed to the introduction and the discussion. MS, HS, and AK critically revised the manuscript. All authors gave 1nal approval for publication.

### Availability of data and material

The software used to run all simulations was C#. NET Framework 4 and the scripts are available at https://github.com/safarvafadar/chemotaxis.

### Additional files

Supplamentary files

- Movie S1: A short clip depicting a digital bacterium in a random walk regime.
- Movie S2: A short clip depicting a digital bacterium motility based on our SML model.
- Movie S3: A short clip depicting a digital bacterium in a non-homogeneous Markov regime.

